# In-utero exposure to estrogen-mimicking bisphenols alters bone mineralization in the offspring

**DOI:** 10.1101/2023.12.27.573412

**Authors:** Saikanth Varma, Archana Molangiri, Sreedhar Mudavath, Rajendran Ananthan, Ajumeera Rajanna, Asim K Duttaroy, Sanjay Basak

**Affiliations:** Molecular Biology Division, ICMR-National Institute of Nutrition, Indian Council of Medical Research, Hyderabad, India; Food Chemistry Division, ICMR-National Institute of Nutrition, Indian Council of Medical Research, Hyderabad, India; Cell Biology Division, ICMR-National Institute of Nutrition, Indian Council of Medical Research, Hyderabad, India; Department of Nutrition, Institute of Basic Medical Sciences, Faculty of Medicine, University of Oslo, Oslo, Norway

**Keywords:** Bisphenols, Osteoblast, Bone mineralization, TGF-β1, Bone morphogenetic proteins

## Abstract

Exposure to plastic-derived estrogen-mimicking endocrine-disrupting bisphenols can have a long-lasting effect on bone health. However, gestational exposure to below tolerable daily intake (TDI) of bisphenol A (BPA) and its substitute, bisphenol S (BPS), on offspring’s bone mineralization is unclear. This study examined the effects of in-utero bisphenol exposure on the growth and bone density of the offspring rats. Pregnant Wistar rats were exposed to BPA and BPS (0.0, 0.4 μg/kg bw) via oral gavage from gestational day 4 to 21. The bone density, IGF-1, osteocalcin, and calcium levels were measured by DEXA, ELISA and AAS, respectively. The bisphenol’s action on canonical BMP signaling was examined in osteoblast SaOS-2 cells. Maternal exposure to bisphenols (BPA and BPS) increased the body weight, bone mineral content, and density in the offspring aged 30 and 90 days (p<0.05). Plasma IGF-1, calcium, osteocalcin, and alkaline phosphatase activities were altered in BPA-exposed offspring (p<0.05). The bisphenols exposure to SaOS-2 cells decreased its viability in a dose-dependent manner and promoted the cell cycle progression of the S/G2-M phase (p<0.05). The expression of BMP1, BMP4, and intracellular signalling mediators SMAD1, SMAD5, and RUNX2 mRNAs was altered upon bisphenol exposure in these cells (p<0.05). The bone mineralization index and expression of extracellular matrix proteins such as ALPL, COL1A1, DMP1, and FN1 were downregulated (p<0.05). Bisphenol co-incubation with noggin decreased TGF-β1 expression, indicating its involvement in bone mineralization. Overall, exposure to bisphenols (BPA and BPS) during gestation altered growth and bone mineralization in the offspring by modulating canonical BMP/ TGF-β1 signalling mediators.

**Highlights:**

- Gestational exposure to low doses of bisphenol increases whole-body BMC and BMD in the offspring.
- In-utero BPA exposure increased plasma IGF-1 and gla-type osteocalcin, a marker of osteoblast activity in the offspring.
- Bisphenol exposure modulates Smad-dependent BMP signaling in the SaOS-2 cells.

## 1. Introduction

Bisphenol A (BPA), an endocrine disruptor, is widely used in the production of polycarbonate plastics and is a component of epoxy resins, employed in the food packaging industry, thermal paper receipts, and dentistry (Schaffert et al., 2021). Due to their ubiquitous presence, pregnant women are continuously exposed to bisphenols from different sources and routes. BPA has been detected in maternal serum, cord blood and fetal tissues (Chou et al., 2011). Potential health risks associated with BPA demanded that the industry replace it with its predominant substitute, bisphenol S (BPS). Bisphenols are xenoestrogen compounds having structural similarity with 17β-estradiol and can bind to estrogen (E2) receptors (ER) with lower affinity than E2 (Kuiper et al., 1998). Unlike adults, fetuses are more susceptible to endocrine programming *in utero* from the bisphenol-exposed mother since they lack protective enzymatic machinery (Basak et al., 2023). Fetal exposure to these endocrine-disrupting chemicals (EDC) can lead to disrupted bone growth (Iwobi and Sparks, 2023) and risks for bone diseases in later life (van Zwol-Janssens et al., 2020).

E2 plays a pivotal role in bone growth and development by modulating bone tissue homeostasis, as evidenced by the presence of ER on osteoblasts, osteocytes and osteoclast cells (Khalid and Krum, 2016). Any disruption in these cells may lead to bone abnormalities such as osteoporosis and osteopetrosis (Lazner et al., 1999). Bone homeostasis is dynamic, where old tissues are removed, and new tissues are formed by maintaining the cellular balance between osteoblast and osteoclast activities. Excess bone tissue resorption in the absence of E2 results in a loss in bone mass, disturbs internal bone architecture and reduces bone strength. EDCs such as BPA and BPS are E2 mimics that can interfere with bone metabolism (Thent et al., 2018). Bisphenol exposure could promote the risk of hormonal imbalances, leading to adverse consequences for the structure and function of bone tissue (Chin et al., 2018). The post-menopausal women with low levels of E2 are more prone to develop osteoporosis, a condition associated with rapid loss of bone mineral density, resulting in fragile bone (Cauley, 2015), indicating potential risks could arise from estrogen-mimicking bisphenols on bone metabolism. Estrogen deficiency induced rat model, resulting in subchondral bone resorption and articular cartilage degeneration (Xu et al., 2019).

As an endocrine disruptor, BPA disturbs bone health by decreasing plasma calcium levels and inhibiting calcitonin secretion (Suzuki et al., 2003). BPA inhibits osteogenic activity and promotes bone resorption (Maduranga Karunarathne et al., 2022). BPA exposure during early development reduces bone stiffness and induces bone marrow fibrosis and chronic inflammation in the offspring (Lind et al., 2019). In addition to estrogen, bone homeostasis is also modulated by IGF-1 (Yakar and Rosen, 2003), ALP (Beertsen and van den Bos, 1992), Osteocalcin (Manolagas, 2020), and others. Previous studies have shown that BPA can affect bone morphology (Prasse et al., 2022) by supersession osteoclastic and osteoblastic activities (Suzuki and Hattori, 2003). Effects of BPA administration showed retardation in the ossification of fetal rats skeleton (Kim et al., 2001). The femoral length of the rats exposed to BPA was significantly higher than that of controls. Fetal exposure to BPA altered femoral diaphyseal BMC, cortical cross-sectional area, and cortical BMC in rat offspring (Lejonklou et al., 2016). BPA also significantly suppressed ALP activities and bone nodule formation in osteoblast precursor cells (Hwang et al., 2013).

Bisphenol mediates its adverse effects on bone homeostasis through various pathways, including transforming growth factor-β (TGFβ)/BMP. The TGF-β is involved in the osteogenic function and bone mass formation (Hu and Chen, 2018) where it binds and activates TGFβR and its downstream effector Smad2/3, which in turn binds to co-Smad 4 and induces transcription of osteogenic differentiation genes that promotes bone formation (Tong et al., 2016). Prenatal BPA exposure impaired bone development and bone mass accumulation via the TGFβ signaling pathway (Wang et al., 2021). Bone morphogenetic protein (BMP) binds and activates the BMP receptors, leading to the activation of Smad 1/5/8, which binds to Smad 4 and translocates to the nucleus to act as a transcription factor and is involved in MSC differentiation and bone formation (Beederman et al., 2013). Prolonged BPA exposure can alter the differentiation potential of stem cells by modulating BMP signaling (Clément et al., 2017). BMP-knock-out mice have increased endogenous bone mass, primarily in trabecular bone, with decreased osteoclastogenesis (Kamiya et al., 2008), indicating the role of BMP in maintaining bone mass.

Despite this evidence, the existing data on the effects of gestational BPA and its substitute, BPS exposure, on the offspring’s bone health are limited. Moreover, the impact of bisphenols on TGF-β and BMP signaling is unknown. The present study investigated the in-utero exposure of below TDI of BPA and BPS on offspring bone health, including whole-body BMC, BMD, plasma IGF-1, ALP, and osteocalcin levels. Additionally, this work evaluated the mechanism of bisphenols’ action on mineralization by examining the TGF-β/BMP signaling mediators in osteoblast SaOS-2 cells.

## 2. Materials and methods

### 2.1 Animal experiment

Healthy age-matched 3-month-old female (n=40) and male (n=20) Wistar rats were procured from the National Institute of Nutrition animal facility. The study design and procedures (#ICMR-NIN/IAEC/02/008/2019) received approval from the Institutional Animal Ethics Committee of ICMR-National Institute of Nutrition, adhering to the guidelines for the care and use of laboratory animals. Rats were housed in two per cage and maintained at 12h light/dark cycle with a temperature of 22°C ± 2°C and relative humidity of 45-55%. Rats were given ad libitum water and a standard rodent chow diet (AIN93M). After acclimatization (7 days), mating was facilitated overnight by introducing male rats (1 male: 2 females). The presence of vaginal plugs confirmed pregnancy and the day was considered as the gestational day (gD) 0. Pregnant rats were divided into three groups, i.e., control (0.0), 0.4 µg/kg bw of BPA and BPS. Bisphenol (BPA & BPS) was dissolved in extra virgin olive oil to get the desired concentrations. Pregnant rats were exposed to bisphenols via oral gavage (0.5 ml/day/rat) from gD 4 to 21. After weaning, the offspring were continued on a similar diet. Blood was collected from the offspring (30 and 90 days) by retro-orbital puncture. The plasma was separated and stored at -80°C for further analysis.

### 2.2 Dual-energy X-ray absorptiometry (DEXA)

DEXA scans were performed to evaluate the rats’ whole-body composition and bone density. Offspring (30 and 90 days) were anesthetized by a cocktail of ketamine and xylazine in a final concentration of 80 and 10 mg/kg body weight, respectively, injected intraperitoneally. A total body scan was performed using DEXA (Discovery, Hologic, Bradford, MA, USA) according to the supplier’s protocol.

### 2.3 Estimation of IGF-1 and Gla-type osteocalcin levels

Plasma insulin-like growth factor-1 (IGF-1) levels were analyzed using ELISA per the manufacturer’s instruction (#E-EL-M3006, Elabsciences, USA). Gla-type osteocalcin levels were measured in plasma and cell culture supernatants. Plasma Gla-type osteocalcin was analyzed by enzyme immunoassay (#MK111, Takara, Japan). The absorbance was read at 450 nm using a microplate reader (Bio-Tek, Powerwave XS). IGF-1 and Gla-type osteocalcin concentrations were expressed as ng/ml.

### 2.4 Determination of calcium by atomic absorption spectrophotometry

Plasma (50 µl) was mixed with diluent buffer containing 0.1 % Lanthanum (III) chloride and 0.1 % suprapur nitric acid, and the final volume was made to 5 ml with diluent buffer. The atomic absorption spectrophotometry (Shimadzu AA-7000) determined plasma calcium concentrations against an ICP multi-element standard solution VIII (Cat #1.09492, Certipur, Supelco). Results were expressed as µg/ml.

### 2.5 Alkaline phosphatase activity

ALP activity was measured in the plasma and cell lysates. Briefly, plasma or cell lysates (15 µl) were made up to 100 µl using water and incubated with 100 µl of 4.4 mM p-nitrophenyl phosphate (PnPP) in 1 mM MgCl_2_. 6H_2_O and 0.05 M glycine pH-10.5, at 37°C for 30 min. After incubation, the reaction was arrested by adding 100 µl of 0.1 N NaOH. Absorbance was read at 405 nm using a microplate reader (Bio-Tek, Powerwave XS). P-nitrophenol (PnP) was used for the standard curve. ALP activity was expressed in IU/L for the plasma samples and nanomoles of p-nitrophenol per min per milligram protein for cell lysates.

### 2.6 Reagents and cell culture

The HTB-85/SaOS-2 osteoblast cell line was obtained by the American Type Culture Collection, USA. L-Ascorbic acid (#A4544), Bisphenol A (#239658), Bisphenol S (#103039), β-Estradiol (#E2758), Methyl thiazolyl diphenyl-tetrazolium bromide (MTT#5655), Potassium phosphate monobasic (#60218), and 4-Nitrophenol (#1048) were obtained from Sigma Aldrich, Germany. Penicillin-streptomycin solution (#15140122), Trypsin-EDTA (#25300062), and Fetal Bovine Serum (#10270106) were obtained from Thermo Fischer Scientific, USA. Dulbecco’s Modified Eagle Medium/High glucose (DMEM) (# SH30249.01) was obtained from Cytiva, HyClone laboratories. Alizarin Red S dye (#60352) and P-nitro phenyl phosphate disodium salt hexahydrate (#90341) were obtained from SRL Chemicals, India. Recombinant human Noggin protein-carrier free (#6057-NG/CF) was obtained from R&D systems.

The SaOS-2 cell was cultured in DMEM containing 25 mM HEPES, 4.5 g/L glucose, and 4 mM L-glutamine supplemented with 10% heat-inactivated fetal bovine serum (FBS) and 1% antibiotics (50 U/ml penicillin and 50 mg/ml streptomycin). Before treatment, cells were serum starved for 24 h, followed by incubation in 2% FBS and media supplemented with 50 µg/ml L-ascorbic acid and 1.8 mM potassium phosphate monobasic (osteogenic culture media) containing either bisphenols or estrogen. Bisphenols (BPA & BPS) and 17β-estradiol (E2) powders were dissolved in absolute alcohol to prepare stock solutions. Stock solutions were diluted with assay media before the treatment of the cells.

### 2.7 Cell viability by MTT assay

SaOS-2 cells were seeded in 96 well plates with a density of 10×10^3^ cells per well using a multichannel pipette. The cells were starved with low FBS (1%) DMEM media for 4 h, followed by assay media (100μl) containing different concentrations of bisphenol A (BPA), bisphenol S (BPS), and 17β-estradiol for 24 h at 37° C in a 5% CO_2_ humidified incubator. Assay media was replaced with MTT (3-[4,5-dimethylthiazol-2-yl]-2,5 diphenyl tetrazolium bromide)-containing media (0.5ng/ml of DPBS) and incubated for 4h. Formazan crystals were dissolved using DMSO. Absorbance was read at 562 nm using a microplate reader (Bio-Tek, Powerwave XS).

### 2.8 Flow cytometry

Cells were trypsinized, harvested in DMEM containing 10% FBS, washed twice with PBS, and fixed in chilled 70% ethanol. Cells were stained with 50µl of 50µg/ml propidium iodide supplemented with 100 µg/ml of RNase enzyme in the dark for 30 min. The cell cycle analysis was performed and analyzed by measuring the amount of propidium iodide-labeled DNA in ethanol-fixed cells. Cell cycle analysis was conducted in a flow cytometer (BD Aria II, BD Biosciences), and ModFit LT Software analyzed the data.

### 2.9 Quantitative real-time PCR

Briefly, cells were lysed using TRIzol reagent (# T9424, Sigma Alrich), and adding chloroform achieved phase separation. RNA was precipitated from the upper aqueous phase using chilled isopropanol. RNA pellet was dissolved in sterile nuclease-free water. DNase 1 treatment was performed to remove traces of contaminating genomic DNA. The quality and quantity were assessed using Nanodrop (ND1000, Thermo Scientific, USA). Total RNA was converted to cDNA using iScript cDNA synthesis kit (#1708891, Bio-Rad, USA). qRT-PCR was performed on Bio-Rad CFX96 real time (qPCR) system using predesigned KiCqstart primers (Sigma) and TBB green (#RR-208A, Takara) pre-mix Ex-Taq II (**Supplementary Table 1**). The Ct values obtained were used to calculate the relative quantification of mRNA expression of genes using the ddCt method.

### 2.10 Immunoblotting

Cells were harvested in radioimmunoprecipitation (RIPA) buffer containing protease inhibitor cocktail and were homogenised using sonication for two cycles for 10 sec each. Cell lysates were centrifuged and separated, and protein concentrations were estimated using the BCA method. Using a wet transfer system, equal protein concentrations were loaded onto SDS-PAGE, resolved, and transferred onto the nitrocellulose membrane (#1620112, Bio-Rad). Membranes were blocked with skimmed milk incubated overnight at 4°C with primary antibody TGF-β1 (#A2124, Abclonal) with a dilution of 1:1000 followed by incubation with HRP conjugate secondary antibody (source) for 90 min at room temperature. Chemiluminescent immunoblot signals were captured in the gel imaging system using a G-box (model) after adding ECL substrate (source). The band intensity was quantified using ImageJ software 1.50i (NIH, Bethesda, MD, USA)

### 2.11 Detection and quantification of osteoblast mineralization by alizarin red S staining

Cells were washed with PBS and fixed in formaldehyde at room temperature for 15 min. After fixation, cells were washed with distilled water, and 40 mM alizarin red S (ARS), pH 4.1 was added, and incubated at room temperature for 20 min with gentle shaking. Unincorporated dye was aspirated, and wells were washed with water to remove the remaining ARS. Plates were stored at -20°C until dye extraction. To quantify staining, 10% acetic acid was added to each well, and the plate was incubated at room temperature with shaking. The monolayer was then scraped, transferred to a microcentrifuge tube, and vortexed. The slurry was centrifuged, and the supernatant was transferred to a new tube. Ammonium hydroxide was added to neutralize the acid. The supernatant was transferred in 96-well plates and read at 405 nm in triplicate using a microplate reader (Bio-Tek, Powerwave XS).

### 2.12 Statistical analysis

Multiple group comparisons were performed using one-way ANOVA in GraphPad Prism v.8. Every experiment was performed independently and repeated multiple times, as mentioned in the text or figure legends. Data was considered statistically significant when the p-value fell less than 0.05. The data were represented as mean ± standard error of the mean.

## 3. Results

### 3.1 Gestational exposure to bisphenols results in increased bone density and mineralization of the offspring

The whole-body composition was measured by DEXA at 30 and 90 days of age following gestational exposure to bisphenols (0.4µg/kg bw/days) in dams. BPA and BPS -exposed offspring showed a significant increase in body weight (BW), body surface area (BSA), bone mineral content (BMC), and bone mineral density (BMD) compared to control at 30 and 90 days of their age (**Fig.1 A-C**). BPA-alternative chemical BPS exhibited similar effects in altering bone density and mineralization in these offspring.

**Fig. 1.**
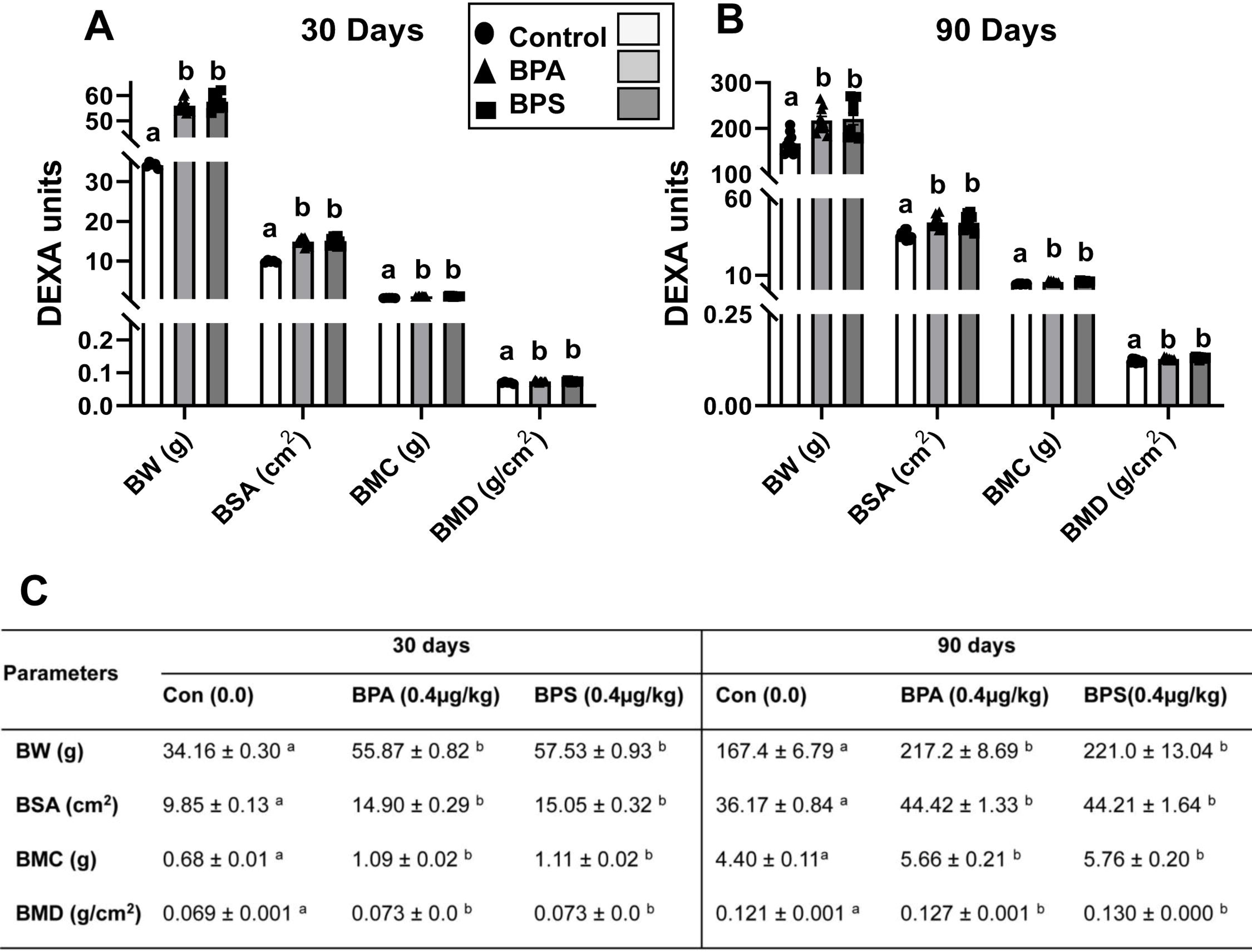
Body mass and bone density of the offspring following *in utero* bisphenols (BPA and BPS) exposure. (A-C) Whole body dual-energy X-ray absorptiometry (DEXA) analysis of the rat offspring at (A) 30 days (B) 90 days of their age. (C) Individual values are expressed as mean ± SEM, n=10/group. Data was analyzed by one-way ANOVA with Tukey’s multiple comparison tests. Values with unlike superscript letters were significantly different p < 0.05 vs control. BW; Body weight, BSA; Body surface area, BMC; Bone mineral content, BMD; Bone mineral density, Con; Control, BPA; Bisphenol A, BPS; Bisphenol S.

### 3.2 In utero, bisphenol exposure dysregulates plasma IGF-1, ALP, and calcium homeostasis in the 90-day offspring

Insulin-like growth factor 1 (IGF-1) levels were significantly increased in the plasma of BPA-exposed offspring (**Fig.2A**), while BPS exposure had no effects. In contrast, the plasma alkaline phosphatase levels increased dramatically in BPS-exposed offspring. Still, not in BPA-exposed rats (**Fig.2B**). Conversely, similar but inverse findings were found in plasma calcium levels, where calcium levels were decreased significantly in BPS-exposed adult rats (**Fig.2C**). The Gla-type osteocalcin levels were significantly increased in the plasma of BPA and BPS-exposed offspring compared to the unexposed control group (**Fig.2D**). Thus, overall data suggests that gestational exposure to bisphenols impaired offspring’s IGF-1, ALP and calcium homeostasis in their adult life.

**Fig. 2.**
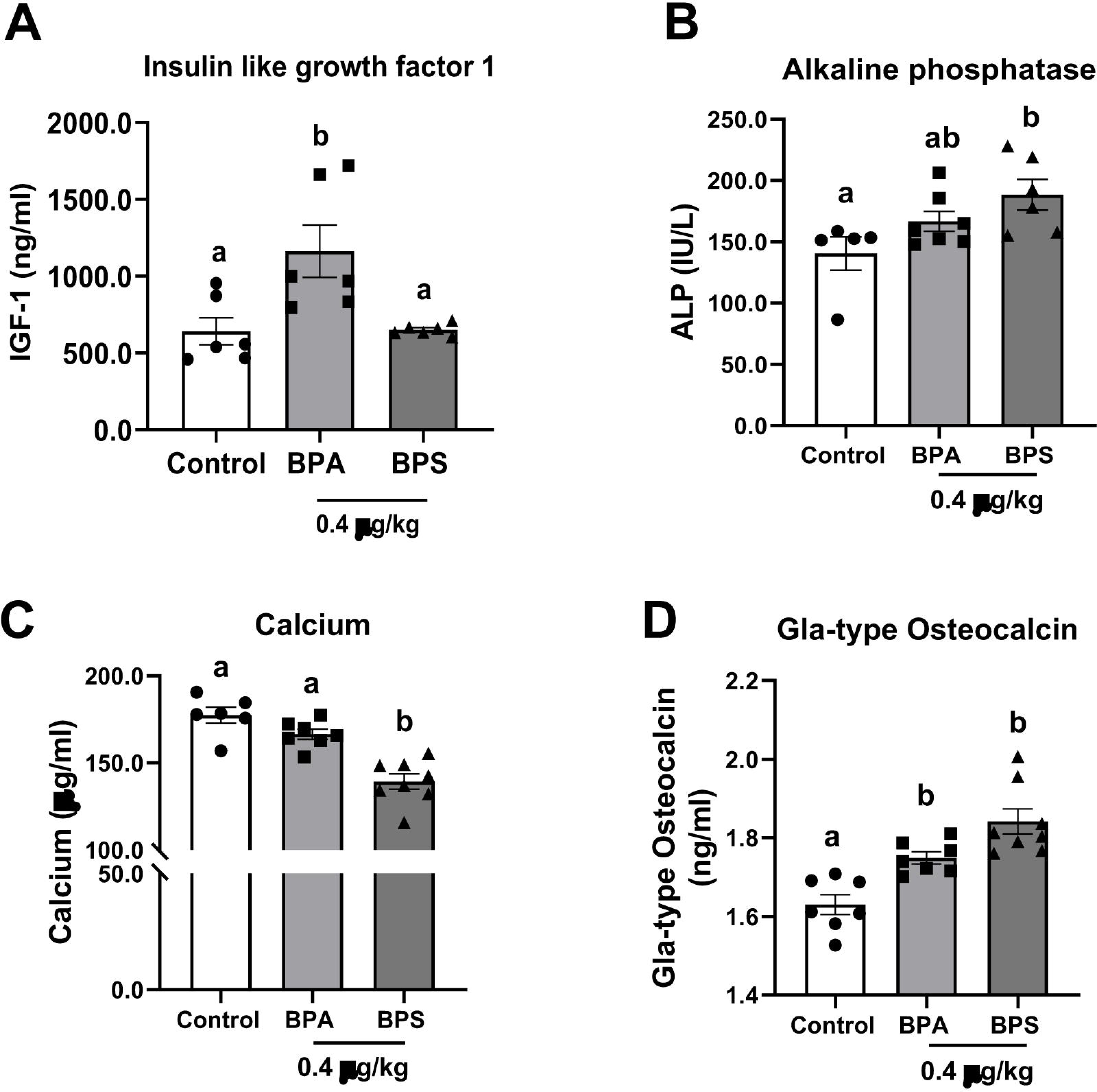
Plasma insulin-like growth factor-1 (IGF-1), alkaline phosphate (ALP), calcium, and Gla-type osteocalcin levels in the offspring (90 days) exposed to bisphenols (BPA and BPS) *in utero*. (A) IGF-1 (ng/ml) level (n=6/group), (B) ALP (IU/L) level (n=5-7/group), (C) Calcium (µg/ml) level (n= 6-8/group), (D) Gla-type osteocalcin (ng/ml) level (n=7-8/group). Data was analyzed by one-way ANOVA with Tukey’s multiple comparison tests and are represented as mean ± SEM. Values with unlike superscript letters were significantly different where p < 0.05 vs control.

### 3.3 Bisphenols exposure decreases the number of viable osteoblasts SaOS-2 cells

To assess the effect of bisphenols (BPA & BPS) on osteoblast viability, we performed a dose-response assay on SaOS-2 cells using an MTT assay. Compared to the control, cell viability significantly decreased when exposed to different concentrations (0.1 to 100 µM) of BPA and BPS for 24 h (**Fig.3A**). At 10µM, viability of SaOS-2 cells was decreased significantly due to bisphenol exposure [control vs. BPA and BPS (%): 99.95 ± 1.82 vs. 82.25 ± 1.88 and 91.05 ± 1.90, n=6]. The 17β-estradiol (E2) was used as a positive control for bisphenols. Cells exposed to E2 varying from 10 to 0.1 µM did not affect cell growth and viability (**Fig.3B**). However, exposure to 100µM estradiol significantly reduced the cell viability (control vs. 17β-estradiol: 99.45 ± 1.82 vs. 53.86 ± 0.89, n=6). These data suggest that BPA has a comparable effect with E2 but a more significant impact on SaOS-2 cells’ viability than BPS at similar concentrations. Cellular expression of estrogen receptor alpha (ESR1) was significantly downregulated in BPA-exposed groups indicating its expression in these cells (**Fig.3C**).

**Fig. 3.**
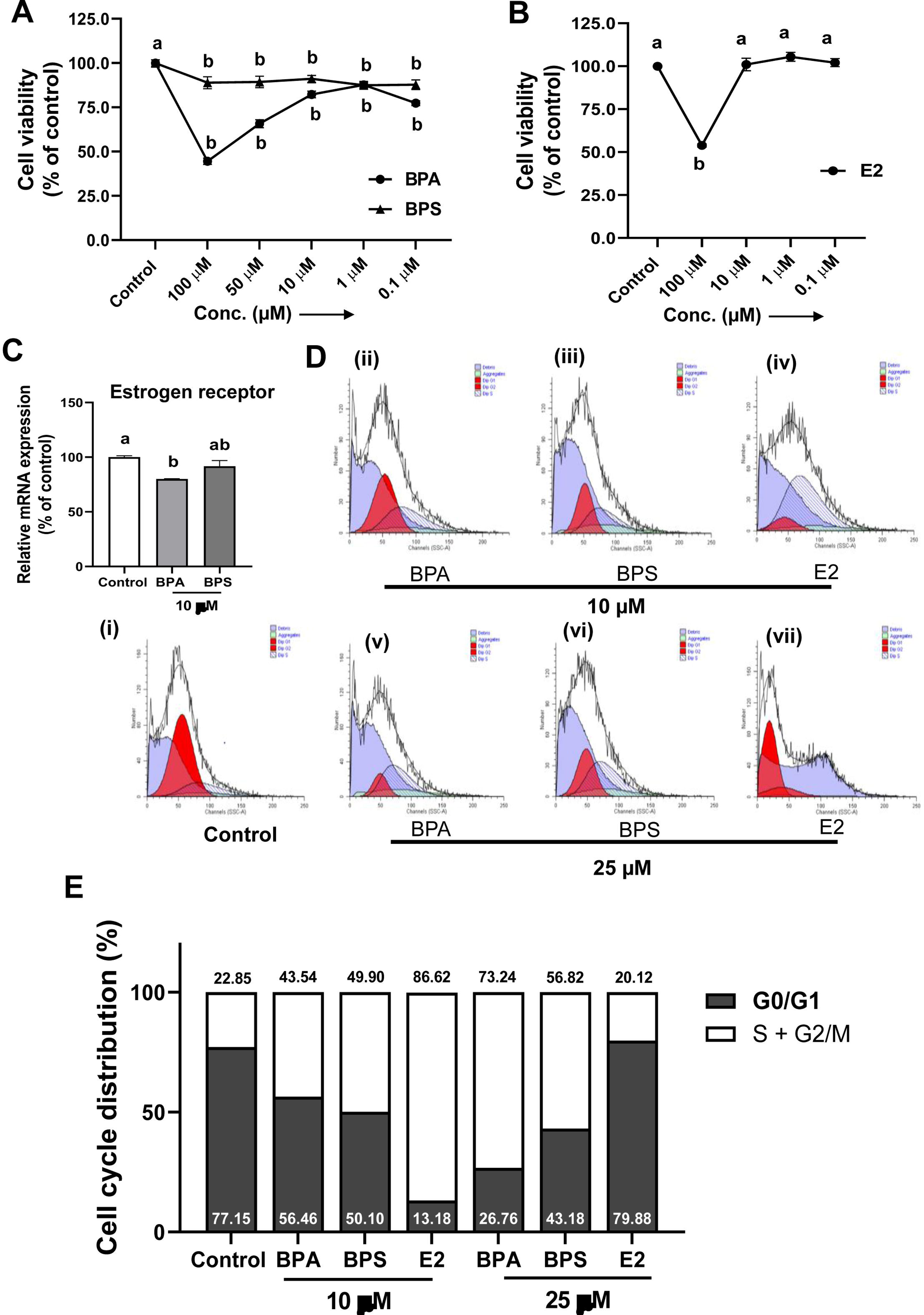
Viability and proliferation of the osteoblast SaOS-2 cells exposed to (A) BPA & BPS (0.1 to100 µM) and (B) E2 (0.1 to100µM) for 24 h. Control cells (vehicle) received 0.1% ethanol. OD values between control and treatment were compared after normalization with blank and expressed as a percentage of control. Cells were exposed to bisphenols (10µM) for 72h in osteogenic culture media as described in the method. (C) The mRNA expression of estrogen receptor was analyzed by qRT-PCR normalized to the endogenous control, GAPDH, calculated according to the ddCt method. Data are represented as mean ± SEM. (D) Analysis of cell cycle progression by flow cytometry. Pictorial graph shows the proportion of cells in different phases of cell cycle after treated with (i) control, (ii, iii, v, vi) bisphenols (10 µM and 25 µM), and (iv and vii) E2 (10 µM and 25 µM) at 72 h. (E) The quantitative expression of cell cycle phase distribution. Data are expressed as mean ± SEM. Groups with unlike letters are significantly different. p < 0.05 vs. control

### 3.4 BPA and BPS exposure increases cell cycle progression by promoting the S-G2/M phase

The effects of bisphenols and estradiol on cell cycle distribution were examined by flow cytometry after treating the cells with BPA, BPS and E2 (10 and 25 µM) for 72 h (**Fig.3D**). Both BPA and BPS significantly increased the cell cycle progression from G0/G1 to S/G2-M phase and decreased the percentage of cells present at the G0/G1 phase in a dose-dependent manner (**Fig.3E**). Like bisphenols, E2 (10 µM) increased the proportion of cells at the S-G2/M phase of the cell cycle compared to the control.

### 3.5 The mRNA expression of bone morphogenetic proteins, and signaling mediators were altered following exposure to bisphenols

The mRNA expression of BMP1 and BMP4 was decreased considerably in BPS (10µM) exposed cells. At the same time, BPA had no effects (**Fig.4A**). In contrast, the exposure to BPA (10µM) significantly increased the BMP2 and BMPR1A mRNA expression. The expression of FGF23 was significantly downregulated in both BPA and BPS-exposed SaOS-2 cells. However, mRNA expression of BMP7, BMPR1β, and BMPR2 was unaffected upon bisphenol treatment (data not presented). Compared to the control, the mRNA expression of transcription factors such as RUNX2, SMAD5, and BMP antagonists such as SOST were significantly upregulated due to exposure to BPA (**Fig.4B**). Conversely, the SMAD1 mRNA expression level was significantly downregulated in BPA-exposed cells but not in the BPS group.

**Fig. 4.**
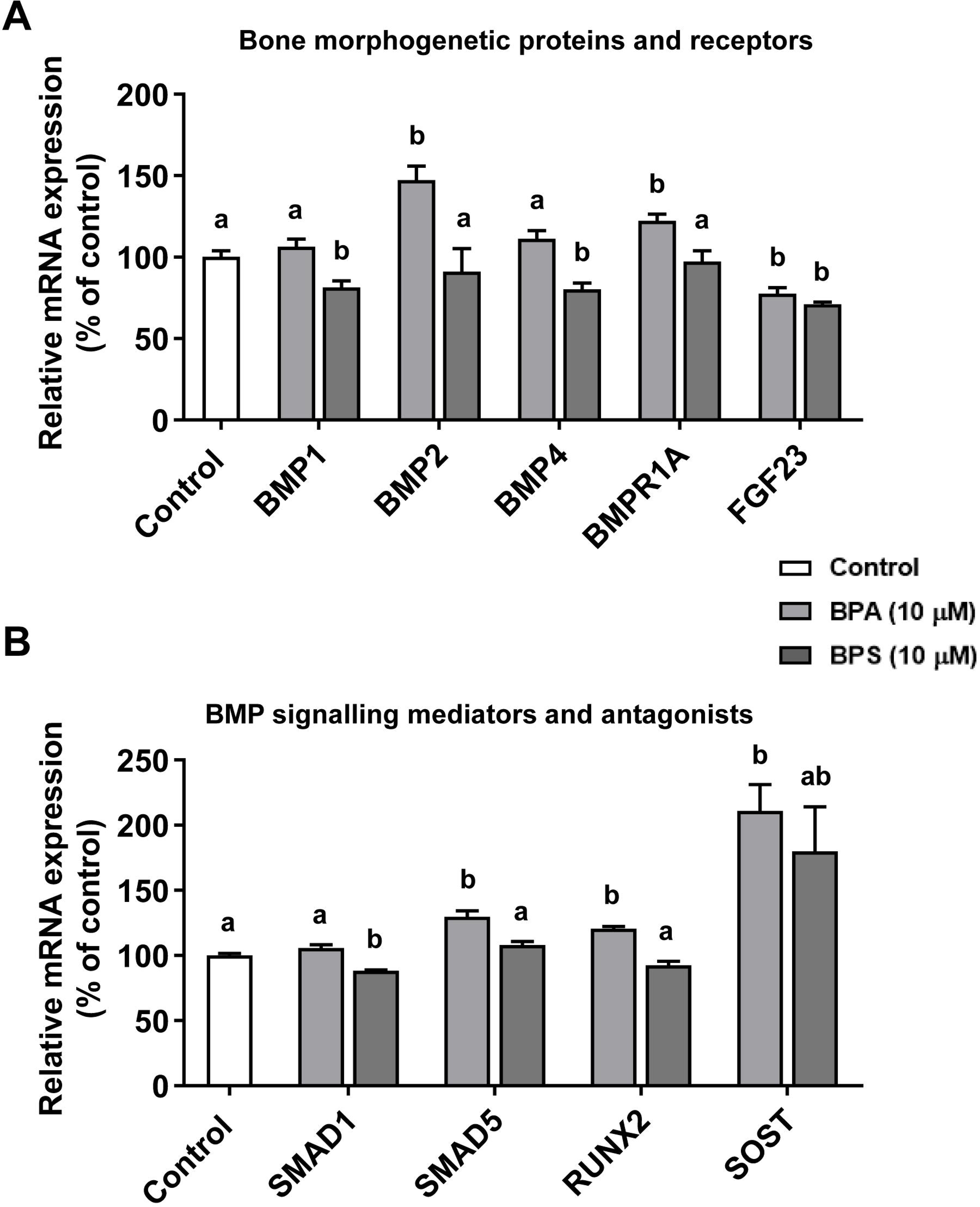
The expression of bone morphogenetic protein (BMP), its receptors and intracellular BMP signaling mediators in the osteoblast SaOS-2 cells following bisphenol exposure. Cells were exposed to bisphenols (10µM) for 72h in osteogenic culture media as described in the method. The mRNA expression of genes was analyzed by qRT-PCR normalized to the endogenous control, GAPDH, calculated according to the ddCt method. (A) Bone morphogenetic proteins and receptors (B) BMP signalling mediators and antagonists. Data are expressed as mean ± SEM (n=4). Groups with unlike letters are significantly different. p < 0.05 vs. control.

### 3.6 In vitro, mineralization and expression of extracellular matrix synthesis genes in SaOS-2 cells were decreased upon bisphenol exposure

BPA and BPS (10µM) significantly decreased calcium nodule formation, as evidenced by a reduction in Alizarin red S levels (**Fig.5A**). Due to intermittent exposure of E2 for 6h of every 48h of culture to SaOS-2, cells increased the calcium nodule formation. The ALP activity was significantly increased in both BPA and BPS-exposed cells. However, the continuous E2 exposure reduced the ALP activity compared to the control group (**Fig.5B**). Expression of several bone mineralization markers was modulated after exposing SaOS-2 cells to BPA and BPS. The mRNA expression of genes associated with extracellular matrix syntheses and bone mineralization, such as COL1A1, ALPL, FN1, and DMP1, were significantly downregulated (**Fig.5C**). However, the expression of PHEX and BGLAP (osteocalcin) increased in both the bisphenol-treated groups. In contrast, expression of SPARC, SP7 and SPP1 mRNAs were unaffected (data not presented). The Gla-type osteocalcin levels in the cell culture supernatants were significantly decreased in the BPS group (**Fig.5D**). In the presence of noggin, a BMP signalling inhibitor, the expression of TGFβ1 was significantly inhibited compared to basal (control) levels (**Fig.5E**). Moreover, coincubation of noggin with BPA and BPS resulted in further inhibition in the expression of TGFβ1 compared to noggin alone. While E2 significantly inhibited the TGF β1 expression independently of noggin. However, BPA and BPS do not affect TGF β1 protein expression alone in the osteoblast SaOS-2 cells. These data suggest that bisphenol alters the expression of TGFβ1 protein with noggin in SaOS-2 cells.

**Fig. 5.**
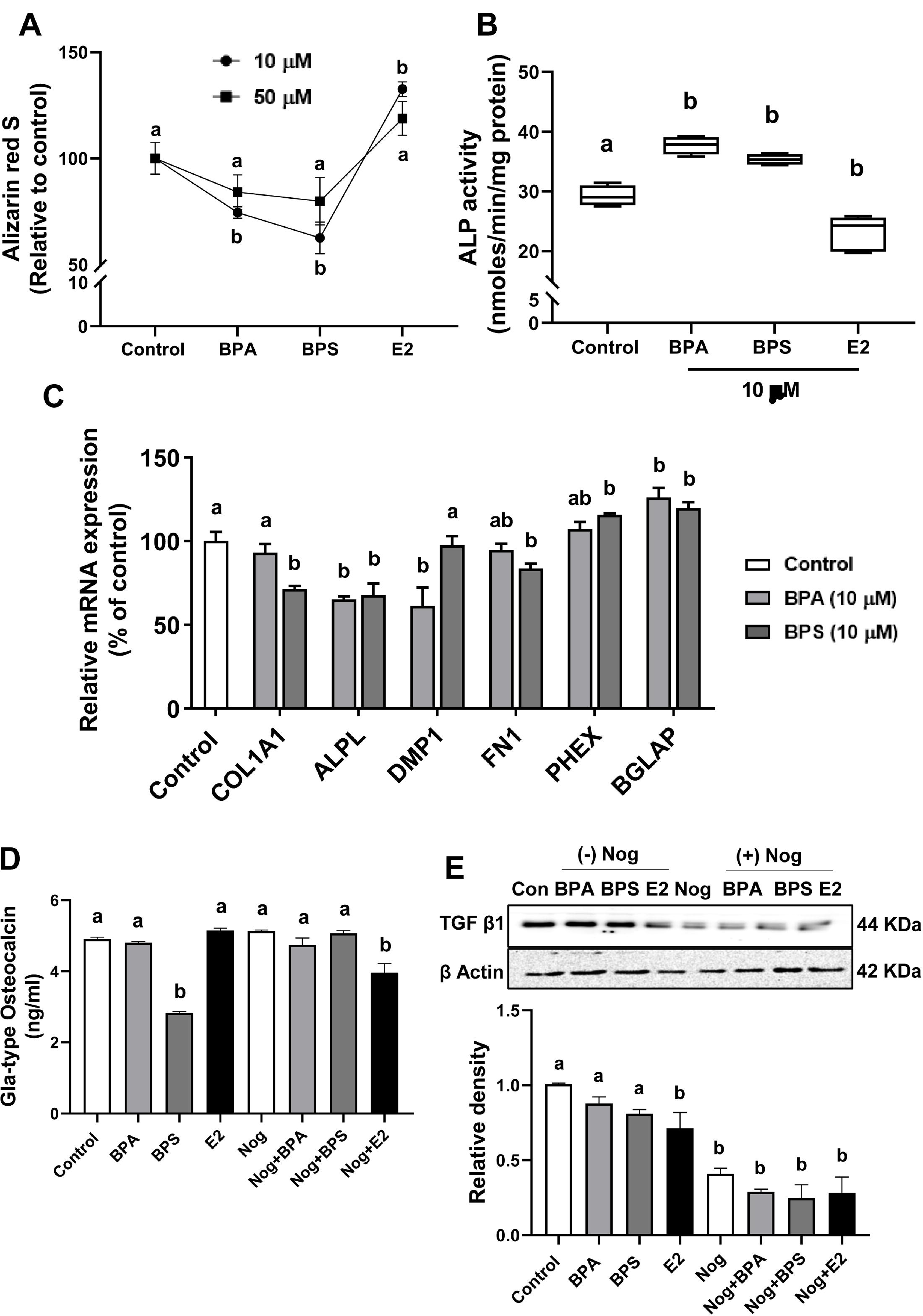
Calcium nodule formation (in vitro mineralization), alkaline phosphatase (ALP), Gla-type osteocalcin levels, and gene expression of bone mineralization and extracellular matrix synthesis markers in osteoblast SaOS-2 cells following bisphenol exposure. (A) Effect of bisphenols on mineralization of the SaOS-2 cells treated with 10-50 µM of BPA, BPS, and E2 for 21 days in osteogenic culture medium as determined by Alizarin red S stain. Intermittent addition was carried out by adding E2 for 6 h of every 48 h of the culture for 21 days. OD values between control and treatment were compared after normalization with blank values and expressed as a percentage of control. (B) In osteogenic culture media, cells were exposed to 10µM bisphenols (BPA and BPS) for 72 h. ALP activity of the SaOS-2 cells after exposed to 10µM bisphenols and 10µM E2 for 72 h (C) mRNA expression of genes involved in bone mineralization and extracellular matrix synthesis was measured by qRT-PCR, normalized with GAPDH, and fold changes in the mRNA expression of genes was calculated according to ddCt method. (D) Gla-type osteocalcin measured by EIA in cell culture supernatants of osteoblast treated with bisphenols (10µM), E2 (10µM) and noggin (1ng/ml). (E) Immunoblots of TGF β1 was performed in cell lysates of SaOS-2 cells. The quantitative bars indicate their relative expression in these cells. Data are expressed as mean ± SEM, n=3-4. Groups with unlike letters are significantly different p < 0.05 vs. control.

## 4. Discussion

For the first time, this study demonstrated that gestational exposure to low-dose BPA (0.4 µg/kg bw/d) and its substitute BPS affected the bone mineralization of the offspring rats. Less than TDI levels of bisphenol exposure during gestation significantly increased the body weight, bone mineral content, and bone mineral density in the young (30d) and adult (90d) rat offspring. Longitudinal changes in bone parameters revealed early fetal exposure to bisphenol might have programmed the development of bone osseous tissue, resulting in excess mineralization in the offspring (Pelch et al., 2012). Moreover, these changes were similar in the BPA and its alternate substitute compound BPS exposure. Additionally, exposure to bisphenols (BPA and BPS) affects cell growth, proliferation, mineralization, and expression of genes involved in the BMP signaling in the osteoblast SaOS-2 cells.

The expression of the estrogen receptor was decreased due to BPA exposure in our previous (Molangiri et al., 2022) and present study, indicating that any disturbances in the hormonal system can affect bone homeostasis (Streicher et al., 2017). Bone, an osseous tissue, secretes and relies on the hormones produced by other tissues and organs (liver and thyroid) to maintain bone homeostasis. During early development, transient alterations in estrogen levels modulate bone turnover through bone cell imprinting, rendering osteoclast activity and increasing bone mass (Migliaccio et al., 2000). Bisphenols, also known as environmental xenoestrogens due to their structural similarity with estrogen, affinity to bind to estrogen receptor (ER) and activate ERs, could mediate their effects by modulating endocrine balance through estrogen activities. Even though BPA has 1000 to 2000-fold less affinity to the ER than 17β-estradiol (E2) (Bolli et al., 2010), the exposure window might be critical for bone formation and development. Like the present observation about increased BMD and BMC in bisphenol-exposed offspring, previous reports with BPA exposure from gD 7 to 22 showed a significantly thicker diaphyseal cortex and a higher total and cortical BMC in the offspring rats (Lejonklou et al., 2016). However, the mechanism of BPA’s modulation of bone mineralization is not understood. In estrogen-deficient knock-out models, BPA exhibited strong estrogenic effects and improved the total BMD in a dose-dependent manner (Toda et al., 2002).

The increased BMD in BPA-exposed offspring observed in this study could be due to increased lean mass (Molangiri et al., 2023) or an excess amount of estrogen produced by the adipose tissue depots (Nguyen et al., 2020). Bisphenol-exposed offspring rats showed similar increases in body weight and bone parameters such as BMC and BMD in Wistar rats fed with a high fat/high sucrose diet, indicating that obesogenic effects on bone mineralization induced by bisphenol are similar to dietary insults (Lecka-Czernik et al., 2015; Malvi et al., 2014; Minematsu et al., 2018). Increased BMD in bisphenol-exposed offspring could be due to adipose hypertrophy (Molangiri et al., 2023) that produces excess estrone (E1), convert it to active form as E2 (Hetemäki et al., 2021). The estrone produced by adipose tissue showed a positive correlation with BMD and relative fat mass, indicating the potential role of estrone in modulating bone mass in BPA-exposed offspring (Suzuki et al., 1995).

The mechanism of BPA’s modulation of bone mineralization could involve several factors, such as estrogen, aromatase and other adipokines. Many in vivo studies have conspicuously shown that BPA exposure can increase the expression and activation of aromatase in different tissues (Castro et al., 2013; Kim et al., 2010). The local production of estrogen by osteoblast and osteoclast cells is governed by aromatase, which catalyses the conversion of C19 androgens to C18 estrogen (Eyre et al., 1998). Aromatase overexpression in osteoblast mice resulted in increased whole-body BMD, and cortical thickness and a reduced number of osteoclasts (Sjögren et al., 2009), indicating that bisphenol exposure might have altered bone mineralization by modulating aromatase expression.

In addition, bisphenols are well known for their obesogenic effects. The increased fat mass, fat percentage, and leptin overexpression in the bisphenol-exposed offspring, observed in our previous study (Molangiri et al., 2023), might have affected bone homeostasis by excess mechanical load. Beyond energy metabolism, leptin stimulates endochondral ossification and osteoblast differentiation (Steppan et al., 2000). Additionally, leptin treatment in leptin-deficient ob/ob mice increases bone formation, BMC, and BMD and inhibits the differentiation of bone marrow mesenchymal stem cells to adipocytes (Hamrick et al., 2005). Thus, increased body weight, fat mass and altered leptin receptors could promote osteoblast activity, thereby increasing the BMC and BMD of the offspring. Overweight and obesity, a metabolic disorder, can increase the mechanical load on the bone, thereby increasing bone mineral density. Osteocytes can sense the changes in the mechanical load and release specific signaling molecules, such as prostaglandins, which can promote bone formation by increasing osteoblast activity (Matsuzaka et al., 2021).

Bisphenols, as EDCs, can alter the hormonal milieu in the body (Zeng et al., 2022). The elevated plasma insulin-like growth factor-1 (IGF-1) observed in our study might have increased BMD and BMC in the BPA-exposed offspring. A positive correlation exists between IGF-1 and total BMD (Rusińska and Chlebna-Sokół, 2006). IGF-1 is a hormone released by the liver that controls bone growth, skeletal maturation, bone mass acquisition, and maintenance of bone health. The increased IGF-1 levels upon BPA exposure could probably enhance the proliferation and differentiation of the osteoblast precursor cells (Crane et al., 2013; Xi et al., 2016). IGF-1 stimulates osteoblast differentiation by activating mTOR signaling and is essential to maintaining bone mass and microarchitecture in mice (Xian et al., 2012). IGF-1-deficient mice exhibited skeletal malformations and delayed mineralization suggesting the role of IGF-1 in bone density during its development (Wang et al., 2006). In contrast to BPA, the IGF-1 levels in the plasma of BPS-exposed offspring remained unchanged. Most IGF-1 are bound (binary and ternary complex), and its bioavailability is modulated by IGF binding proteins (IGFBPs), especially the predominant IGFBP-3. IGF-1 to IGFBP-3 molar ratio is a surrogate marker of biologically active IGF-1 (Haj-Ahmad et al., 2022). Despite increased body weight and bone density in the BPS-exposed offspring, unaltered IGF-1 levels could be due to reduced distribution of circulating IGF-1 between ternary complex (IGF1/IGFBP3/ALS), binary complex (IGF1/IGFBP3), and free forms.

We found increased ALP activity in the offspring’s plasma exposed to bisphenols, indicating an alteration in the osteoblastic activities due to exposure, as reported earlier (Hassan et al., 2012; Pal et al., 2017). Increased ALP levels enhance osteoblastic activity and are key factors needed for mineralization and bone formation (Vimalraj, 2020). In our study, gestational exposure to bisphenols increased the Gla-type osteocalcin in the plasma of adult offspring, suggesting elevated osteocalcin levels in these rats could have mediated increased bone formation. The osteocalcin (OC) is a vitamin K-dependent calcium-binding protein with carboxylated glutamic acid residues, which mediates strong binding of OC to hydroxyapatite. The OC level can indicate the overall activity of cells operating in bone formation (Deftos et al., 1982). On the contrary, the plasma calcium levels were decreased in bisphenol-exposed offspring, indicating the circulatory calcium was exhausted for the increased bone mineralization.

We adopted an *in vitro* model of osteoblast SaOS-2 cells to understand the mechanism of bisphenol exposure on mineralization. Present data shows that the viability of SaOS-2 cells was inhibited with increasing concentration of bisphenols. However, at similar concentrations, BPA showed more inhibitory effects than BPS. The potential difference in cell viability between BPA and BPS could be due to their structural differences caused by replacing the central carbon atoms with two methyl groups with sulphonyl groups in BPS (Ben-Jonathan and Hugo, 2016). The concentration of bisphenols used in the study was similar to that used in osteoblast-like cells, including MG-63 and MC-3T3E1 (Hwang et al., 2013; Kanno et al., 2004). Furthermore, exposure to BPA and BPS in osteogenic culture increases cell cycle progression into the S/G2-M phase of cell division, as evidenced by BPA’s similar actions in other cells (Deng et al., 2021; Lin et al., 2019).

The BPA exposure significantly dysregulated the expression of BMPs (BMP1, BMP2, and BMP4), receptors (BMPR1A), and intracellular signaling mediators (Smads) in our study, indicating its effects on osteoblast differentiation could be mediated by modulating transcriptional regulation of BMP signaling mediators. The impact of bisphenols on signaling pathways involved in mineralization and BMP signaling mediators is not clearly understood. Bone morphogenetic protein BMP1 and BMP4 are the members of the TGF-β superfamily involved in the processing of extracellular matrix proteins and maintaining bone homeostasis (Wang et al., 2014). BMPs bind to BMPR1A and BMPR1B, activating several downstream signaling pathways through smad proteins that enhance the expression of osteogenic genes. BMP2 increases osteocalcin expression and induces bone formation (Noël et al., 2004) by stimulating osteoblast differentiation (Ogasawara et al., 2004).

Present data showed that BPA exposure significantly dysregulated the mRNA expression of BMPR1A and FGFR3 receptors. However, the mechanism of BPA’s action in modulating transcriptional regulation is unclear. Due to structural similarity with steroid hormone, bisphenols can quickly diffuse across the cell membrane, and can imitate or bind to membrane or cytosolic receptors, leading to altered gene expression response (Basak et al., 2023). In our work, the altered expression of regulatory Smads such as Smad1 and Smad5 levels in the BPA-exposed osteoblast cells indicates that modulation of Smad-mediated BMP signaling could affect bone matrix synthesis, mineralization, and osteoblast function (Zou et al., 2021).

Despite an increase in the expression of osteogenic transcription factor RUNX2, the genes involved in the synthesis of extracellular matrix synthesis (COL1A1, ALPL, DMP1, and FN1) were downregulated in our study, thereby decreasing in vitro mineralization in osteoblast cells. Interestingly, bisphenol exposure increased the sclerostin (SOST) mRNA expression in osteoblast cells, indicating that BPA and BPS might inhibit BMP signaling by upregulating SOST, thereby decreasing the calcium nodule formation. SOST is secreted by terminally differentiated osteocytes embedded within the newly formed bone matrix (Mullen et al., 2013) and it has catabolic effects on bone and inhibits the BMP signaling in SaOS-2 cells (Krause et al., 2010) indicating that bisphenol might have altered bone mineralization by modulating the BMP signaling in osteoblast cells.

TGF-β and BMP signaling is crucial in embryonic skeletal development, bone formation, and homeostasis. Moreover, to understand the effect of bisphenol on TGF-β1 protein expression, we co-incubated the noggin with bisphenols. Co-treatment of bisphenols and noggin significantly decreased TGF-β1 expression compared to control and noggin alone. Noggin is an antagonist of BMP signaling and inhibits it by binding to BMP2, BMP4, and BMP7 (Zimmerman et al., 1996). A recent study revealed that noggin also acts as a potent antagonist for TGF-β1 (Wen et al., 2021).

The TGF-β1 deficient mice showed a significant loss of trabecular bone density and reduced osteoblast number on the bone surface (Tang et al., 2009) suggesting essential role of TGF-β1 for maintaining bone mass. Changes in the expression of mineralization markers such as COL1A1, ALPL, and DMP1 indicated that bisphenol exposure might have affected extracellular matrix synthesis and calcium nodule formation.

Prolonged exposure (21d) to bisphenols in an osteogenic medium inhibited the calcium module formation in the SaOS-2 cells observed in our study, indicating its adverse effects on mineralization processes. Many studies have reported decreased mineralization due to bisphenol exposure in different cells, such as MG-63 (Maduranga Karunarathne et al., 2022), MC3T3-E1 (Hwang et al., 2013), primary human osteoblast (García-Recio et al., 2022; García-Recio et al., 2023). The potential discrepancies between in vitro (calcium nodule formation) and in-vivo mineralization observed in our study could be due to the involvement of system’s biological response, including hormonal regulation, rather than the cell culture system. These data suggest that BPA and BPS modulate in-vitro mineralization by TGF-β and BMP signaling via the TGF-β/BMP/Smad pathway.

The study has several limitations, including the lack of analysis of mineralization mediators in bone tissue. Additionally, in this animal study, we have not focussed on the sex-specific effects of bisphenol on BMC and BMD, as the outcome of gestational exposure to BPA and BPS remained similar in both genders of the offspring.

## 5. Conclusions

Our study, for the first time, demonstrated that prenatal bisphenol exposure affected bone mineralization in the Wistar rat offspring and modulated the TGF-β/BMP signaling in the SaOS-2 osteoblast cells. Different reasons can be attributed to altered BMC and BMD, including altered IGF-1 levels and increased body weight coupled with high-fat mass and percentage, which can increase mechanical load and excess adipokines and non-gonadal estrone hormone production by adipose tissue. Our data suggests that less than TDI level exposure of BPA (0.4 µg/kg bw/day) to pregnant dams severely impacted the body weight, BMC, and BMD in the offspring fed with a standard chow diet. The effects were similar in its predominant BPA substitute, “BPS.” Moreover, the *in vitro* findings revealed that bisphenols could mediate their action on mineralization through TGF-β/BMP signaling in osteoblast cells.

Present preclinical data suggest that in-utero exposure to estrogen-mimicking BPA and its substitute BPS altered bone mineralization, which might have a long-lasting effect on bone health in the offspring. However, a similar risk of exposure to bisphenols on bone health must be examined in humans in a well-controlled clinical trial.

## Abbreviations

AAS: Atomic absorption spectrometry
ALP: Alkaline phosphatase
ARS: Alizarin Red S
BPA: Bisphenol A
BPS: Bisphenol S
BMC: Bone mineral content
BMD: Bone mineral density
BMP: Bone morphogenetic protein
bw: Body weight
BMPR1A: Bone morphogenetic protein receptor 1A
BMPR1β: Bone morphogenetic protein receptor 1β
BMPR2: Bone morphogenetic protein receptor 2
COL1A1: Collagen type 1 alpha 1
DEXA: Dual energy x-ray absorptiometry
DMEM: Dulbecco’s modified eagle medium
DMP1: Dentin matrix acidic phosphoprotein 1
ER: Estrogen receptor
E2: 17 β-estradiol
EDCs: Endocrine disrupting chemicals
ESR1: Estrogen receptor alpha
FBS: Fetal bovine serum
FN1: Fibronectin 1
gD: Gestational day
IGF-1: Insulin like growth factor 1
MSCs: Mesenchymal stem cells
MTT: Methyl thiazolyl diphenyl-tetrazolium bromide
PHEX: Phosphate Regulating Endopeptidase X-Linked
PnP: P-nitrophenol
PnPP: P-nitrophenyl phosphatase
RIPA: Radioimmunoprecipitation buffer
RUNX2: Runt-related transcription factor 2
SPARC: Secreted protein acidic and cysteine rich
SP7: S7 transcription factor
SPP1: Secreted phosphoprotein 1
SOST: Sclerostin
TGF-β1: Transforming growth factor beta 1
TGFβR: Transforming growth factor receptor

## Authors’ contribution

SV conducted the animal trial, sample and data collection, performed major laboratory experiments, wrote the draft, data curation, data analysis; AM conducted the animal trial, sample, and data collection and performed key experiment. SM, RA, and AR performed key laboratory experiment; AKD provided critical comments and reviewed the manuscript. SB involved in fund acquisition, conceptualized, supervised, administered, drafted, interpreted, and finalized the manuscript.

---

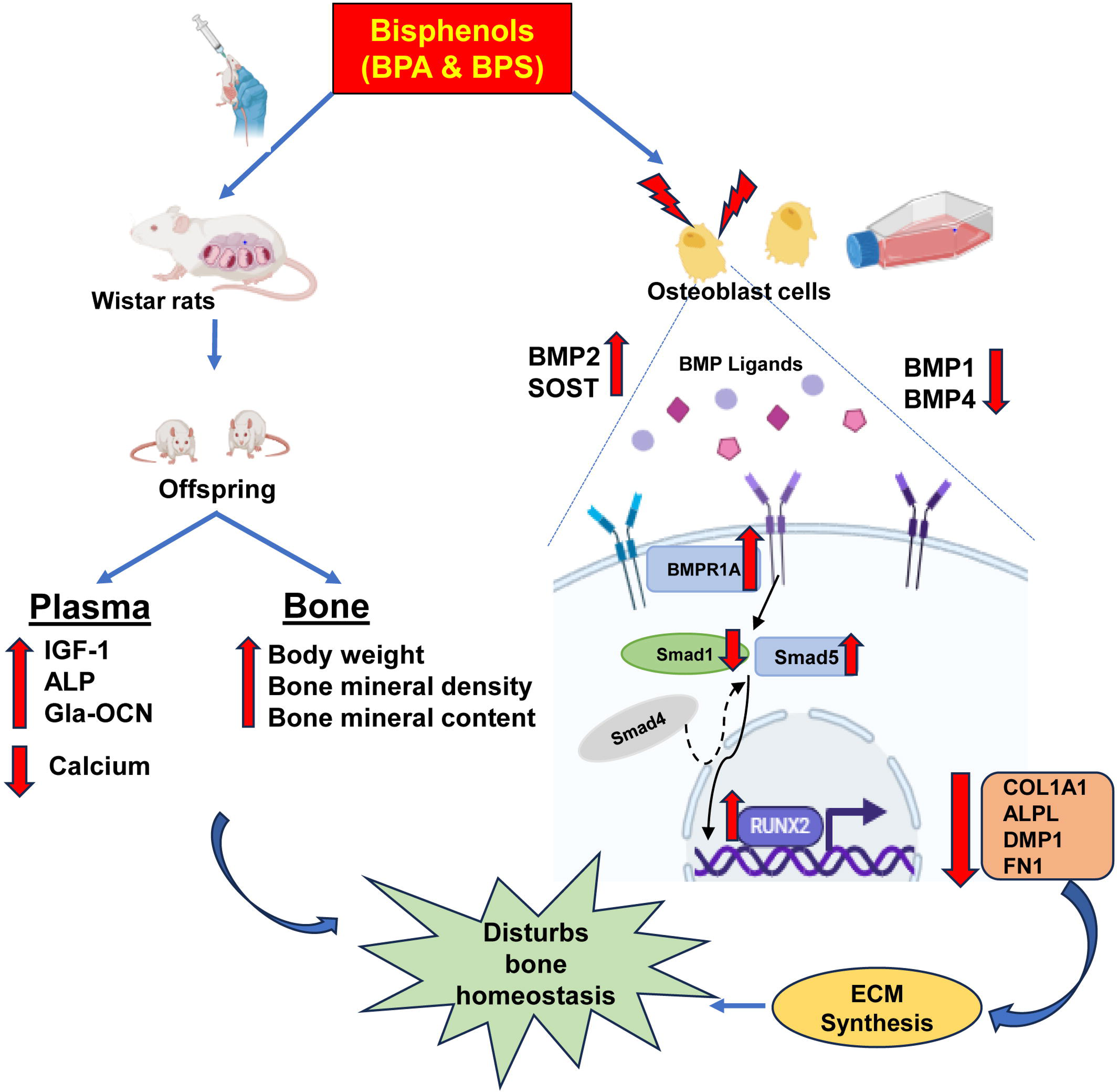

