## Supplementary Table 1 for "In-utero exposure to estrogen-mimicking bisphenols alters bone mineralization in the offspring"

**Table S1**: Predesigned SYBR green I human primers and corresponding genes used for the mRNA expression analyses

| **Sl.**  **no.** | **Primer ID** | **Gene symbol** | **Gene ID** | **Gene name** | **Nucleotide sequences (5’-3’)** | **Ref_seqID** |
| --- | --- | --- | --- | --- | --- | --- |
| 1 | H1_ALPL | ALPL | 249 | Alkaline phosphatase | F 5'- TCTTCACATTTGGTGGATAC-3'  R 5'- ATGGAGACATTCTCTCGTTC-3' | NM_000478.6 |
| 2 | H1_SPP1 | SPP1 | 6696 | Secreted phosphoprotein 1 / osteopontin | F 5'- GACCAAGGAAAACTCACTAC-3'  R 5'- CTGTTTAACTGGTATGGCAC-3' | NM_001040058.2 |
| 3 | H1_BMP2 | BMP2 | 650 | Bone morphogenetic protein 2 | F 5'- TCCACCATGAAGAATCTTTG-3'  R 5'- TAATTCGGTGATGGAAACTG-3' | NM_001200.4 |
| 4 | H1_BMP4 | BMP4 | 652 | Bone morphogenetic protein 4 | F 5'- AGAACATCTGGAGAACATCC-3'  R 5'- AATGTTTATACGGTGGAAGC-3' | NM_001202.6 |
| 5 | H1_BMP7 | BMP7 | 655 | Bone morphogenetic protein 7 | F 5'- TGGTCCACTTCATCAACC-3'  R 5'- TTCTGTATTTCTTCAGGATGAC-3' | NM_001719.3 |
| 6 | H1_BMPR1A | BMPR1A | 657 | Bone morphogenetic protein receptor type 1A | F 5'- CATACTTGGTTTCATAGCGG-3'  R 5'- ATAAGCCAATTTAAGCAGGG-3' | NM_001406559.1 |
| 7 | H1_BMPR1B | BMPR1B | 658 | Bone morphogenetic protein receptor type 1B | F 5'- CATTTTGGGTTTCATTGCTG-3'  R 5'- TGACAGAAGAGTAGGCTAAC-3' | NM_001256793.2 |
| 8 | H1_BMPR2 | BMPR2 | 659 | Bone morphogenetic protein receptor 2 | F 5'- AACAACAGCAATCCATGTTC-3'  R 5'- TATCTFTATACTGCTGCCATC-3' | NM_001204.7 |
| 9 | H1_SMAD1 | SMAD1 | 4086 | SMAD family member 1 | F 5'- GGCATATTGGAAAAGGAGTTC-3'  R 5'- AGATGCTACTGTCACTAAGG-3' | NM_005900.3 |
| 10 | H1_SMAD5 | SMAD5 | 4090 | SMAD family member 5 | F 5'- CCAGTCTTACCTCCAGTATTAG-3'  R 5'- TCCTAAACTGAACCAGAAGG-3' | NM_005903.7 |
| 11 | H1_PHEX | PHEX | 5251 | Phosphate regulating endopeptidase X-linked | F 5'- CTATCAAGTTTTCAGAAGCCG-3'  R 5'- CTTAGCCAGAAGAAATCAGAC-3' | NM_000444.6 |
| 12 | H1_SOST | SOST | 50964 | Sclerostin | F 5'- GAACAACAAGACCATGAACC-3'  R 5'- TACTCGGACACGTCTTTG-3' | NM_025237.3 |
| 13 | H1_ESR1 | ESR1 | 2099 | Estrogen receptor 1 | F 5'- GGAGTGTACACATTTCTGTC-3'  R 5'- CAAAGTGTCTGTGATCTTGTC-3' | NM_000125.4 |
| 14 | H1_SPARC | SPARC | 6678 | Secreted protein acidic and cysteine rich | F 5'- AGTATGTGTAACAGGAGGAC-3'  R 5'- AATGTTGCTAGTGTGATTGG-3' | NM_003118.4 |
| 15 | H1_COL1A1 | COL1A1 | 1277 | Collagen type 1 alpha 1 chaain | F 5'- GCTATGATGAGAAATCAACCG-3'  R 5'- TCATCTCCATTCTTTCCAGG-3' | NM_000088.4 |
| 16 | H1_BGLAP | BGLAP | 632 | Bone gamma carboxyglutamate protein | F 5'- TTCTTTCCTCTTCCCCTTG-3'  R 5'- CCTCTTCTGGAGTTTATTTGG-3' | NM_199173.6 |
| 17 | H1_SP7 | SP7 | 121340 | SP7 transcription factor | F 5'- TGAGGAGGAAGTTCACTATG-3'  R 5'- CATTAGTGCTTGTAAAGGGG-3' | NM_001173467.3 |
| 18 | H1_RUNX2 | RUNX2 | 860 | RUNX family transcription factor 2 | F 5'- AAGCTTGATGACTCTAAACC-3'  R 5'- TCTGTAATCTGACTCTGTCC-3' | NM_001024630.4 |
| 19 | H1_FN1 | FN1 | 2335 | Fibronectin 1 | F 5'- CCATAGCTGAGAAGTGTTTTG-3'  R 5'- CAAGTACAATCTACCATCATCC-3' | NM_212482.4 |
| 20 | H1_BMP1 | BMP1 | 649 | Bone morphogenetic protein 1 | F 5'- GATGTGAAAAAGGACTATGGC-3'  R 5'- AATCTCAAAGGACTGGAATG-3' | NM_001199.4 |
| 21 | H1_DMP1 | DMP1 | 1758 | Dentin matrix acidic phosphoprotein 1 | F 5'- CAACTATGAAGATCAGCATCC-3'  R 5'- CTTCCATTCTTCAGAATCCTC-3' | NM_004407.4 |
| 22 | H1_FGFR3 | FGFR3 | 2261 | Fibroblast growth factor recepptor 3 | F 5'- GAAGATGCTGAAAGACGATG-3'  R 5'- GCAGGTTGATGATGTTTTTG-3' | NM_000142.5 |
| 23 | H1_GAPDH | GAPDH | 2597 | Glyceraldheyde 3 phosphate dehydrogenase | F 5'- ACAACTTTGTCAAGCTCATTTCC-3'  F 5'- GATAGGGCCTCTCTTGCTCA-3' | NM_002046.7 |
